# Isolation, characterization and antibiogram of *Bacillus cereus* from milk products

**DOI:** 10.1101/2024.04.28.591532

**Authors:** Pramod Yadav, Rajesh Khurana, Manesh Kumar, Ritu Yadav, Rinku

**Affiliations:** Department of Veterinary Public Health and Epidemiology, Lala Lajpat Rai University of Veterinary and Animal Sciences

## Abstract

This study, conducted within the Department of Veterinary Public Health and Epidemiology (VPHE) at Lala Lajpat Rai University of Veterinary and Animal Sciences (LUVAS), undertook a comprehensive investigation into the prevalence, identification, characterization, and antibiotic resistance patterns of Bacillus cereus in milk products from various regions of Haryana, India.

Using a systematic sampling strategy, eight tehsils spanning two agroclimatic zones were selected for sample collection. A total of 200 samples were obtained from randomly selected shops within these tehsils. Each sample underwent pre-enrichment in Brain Heart Infusion (BHI) broth with a 1:10 dilution to facilitate the growth of any existing *B. cereus* contaminants. Following pre-enrichment, streaking on selective PEMBA agar plates was performed for the isolation of *B. cereus* colonies. The presence of presumptive *B. cereus* colonies was confirmed through a series of biochemical tests, including gram staining, nitrate reduction, oxidase, indole, methyl red reduction, Voges Praskaur, and catalase tests.

Molecular analysis using the *16SrRNA* gene confirmed the presence of 88 positive *B. cereus* isolates. Further characterization involved the differentiation of *B. cereus* from Bacillus thuringiensis via the cry2 gene. Additionally, the study assessed the presence of virulence-associated genes, identifying *gyrB*, cytk, *hblA*, and *nheA* genes in the isolated strains.

Geographical variation in *B. cereus* prevalence was observed, with higher rates detected in certain tehsils such as Kalka (68%) and Panchkula (60%). Antibiotic sensitivity testing using Tryptone Soya Agar (TSA) broth for enrichment and Muller Hinton Agar (MHA) plates with 14 antibiotic discs revealed widespread resistance among the isolates. Notably, all isolates exhibited resistance to Ampicillin, Cefazolin, Methicillin, Polymyxin-B, and Penicillin, while displaying maximum sensitivity to Amikacin, Gentamycin, Levofloxacin, and Meropenem. Alarmingly, all isolates displayed multiple drug resistance (MDR), indicating resistance to three or more classes of antibiotics.

These findings underscore the urgent need for stringent food safety measures and surveillance protocols in dairy production and distribution. Effective interventions are imperative to mitigate the risk of *B. cereus* contamination and combat antibiotic resistance in milk products. Collaboration with the VPHE department at LUVAS and continued research efforts are essential to address emerging challenges and uphold food safety standards effectively.

## Introduction

Milk is defined as the physiological secretion of the mammary gland of mammals. Milk is necessary for young ones of all mammalian species because it is a primary source of nutrition for them before they are able to digest other types of food. Man has recognized all along the value of milk and milk products as food not only for the young but also for the adults (Nickerson, 1995). It is a healthy and nutritious daily drink that is consumed all over the world and by the people of all ages. Milk provides essential nutrients and also constitutes an important source of dietary energy, high quality proteins and fats.

India is the largest milk-producing nation with estimated production of 230.6 million tonnes with per capita availability of 459 gms/day in 2022–2023: its share in global milk production is 24% (NDDB, 2023). Dairy sector in India plays an important role in socio-economic development and provides livelihood to millions of homes in villages. Consumption of milk and dairy products is deeply rooted in Indian tradition as an important part of daily diet, essential items during rituals and festivals. Changing life style, rising income and urbanisation have affected consumption patterns and increased the demand for more value-added dairy products. Various cooperatives and private sector dairies are producing more dairy products like ghee (clarified butter), butter, yoghurt, khoa, milk powder, paneer (cottage cheese), ice cream, cheese and ethnic sweets to meet this demand. With an increasing demand of dairy products, the need of extended refrigerated storage of raw milk before processing and the application of higher pasteurisation temperatures for prolonged shelf-life requirements have enhanced the importance of thermoduric microorganisms

The incidence of foodborne illnesses has increased globally, and it becomes more important in developing countries where food products are exposed to contaminated environments in food processing industries and temperature abuse during transportation and storage at retail outlets (WHO, 2007). Bacillus cereus (*B. cereus*) is an important safety and shelf-life concern in dairy industry. It is associated with foodborne outbreaks by producing enterotoxins (Anderson Borge *et al*., 2001) and is also responsible for decrease in the organoleptic quality of milk and dairy products by causing spoilage, like sweet curdling and bitterness of milk (Chen *et al*., 2003).

B. *cereus* has been distinguished as a cause of food poisoning since 1955. There are only a few outbreaks per year reported by CDC. Between 1972 and 1986, 52 outbreaks of food-borne illness related to *B. cereus* were reported to the CDC (in 2003, there were two), but this represents only 2% of the total cases which have occurred during these periods. Since, it is not a reportable disease, it usually goes undiagnosed. *B. cereus* causes two types of food-borne illnesses. Emetic type, which is characterized by nausea, vomiting and abdominal cramps within 1 to 6 hours of infection. It resembles *Staphylococcus aureus* (staph) food poisoning symptoms. The diarrheal type is manifested primarily by abdominal cramps and diarrhea has an incubation period of 8 to 16 hours. It may be a small volume or profuse and watery diarrhea. This resembles food poisoning caused by *Clostridium perfringens*. In both types, the illness usually lasts less than 24 hours after onset. In a few cases symptoms may last longer. The short-emetic form is caused by a preformed, heat-stable emetic toxin, ETE. The long-diarrheal form of illness is mediated by the heat-labile diarrheagenic enterotoxin Nhe and/or hemolytic enterotoxin HBL, which causes intestinal fluid secretion, probably by a number of mechanisms, including pore formation and activation of adenylate cyclase enzymes.(Bacillus Food Poisoning. Cambridge City Council.

*B. cereus* group occupies an important position. This group consists of *B. thuringiensis, B. mycoides, B. pseudomycoides, B. anthrasis* and *B. cereus*. (Ankolekar *et al*., 2009). *B. cereus* is a Gram positive, facultative anaerobic, spore forming, motile bacterium (Tallent *et al*., 2012), which is widely distributed in nature and contaminates almost every agricultural commodity (Khudor *et al*., 2012). It is an opportunistic pathogen widely spread in the environment and may provoke harmful food borne illness (Jessberger *et al*., 2019) as well as other infections, such as wound infections, bloodstream infection, umbilical cord infection in neonates, and respiratory tract infections, etc. (Bratcher *et al*., 2017; Prod’hom *et al*., 2017). It is considered as an important enterotoxigenic food borne pathogen causing diarrhoeal and emetic food poisoning (Johnson *et a*l., 1984; Kramer and Gilbert, 1989). It has been reported that the bacterium is isolated from numerous foods, including dairy products, eggs and meat (Kramer and Gilbert, 1989; Ombui *et al*., 2008). *B. cereus* can grow in maximum foods at a pH above 4.5 and temperatures above 4°C (Reyes *et al*., 2001; Svensson *et al*., 2007). *B. cereus* associated food-borne illness occurs as two distinct intoxication syndromes namely emetic and diarrhoeal (Haddaji *et al*., 2022). Apart from gastroenteritis, *B. cereus* has a role in causing variety of non-gastro intestinal tract infections such as meningitis, endopthalmitis, endocarditis, periodontitis, osteomyelitis, wound infections and septicaemia in humans (Schoeni and Wong, 2005). Public health significance of its presence in milk and milk products is due to its heat resistance and potential pathogenic character.

The importance of *B. cereus* in the dairy industry has been recognized since 1938, when the occurrence of bitty cream was recorded. However, *B. cereus* food poisoning was first described in 1950 following consumption of contaminated vanilla sauce (Granum and Lund, 1997). Since 1950, increasing awareness and recognition of *B. cereus* associated illness has resulted in a substantial increase in the number of reports of this type of food poisoning. In India, presence of this organism has been reported in milk (Altaf *et al*., 2015; Chang *et al*., 2021) and other foods of animal origin (Suthar *et al*., 2019).

Antibiotic treatment is still the primary means to eradicate pathogenic *B. cereus*, if this bacterium causes serious infections, but the rapid expansion of antibiotic resistance of *B. cereus* has led to growing difficulty in practical treatment and requires the antibiotic use to be strictly controlled (Yu *et al*., 2019; Yu *et al*., 2020). Thus, it is urgent to develop alternative methods to control pathogenic *B. cereus* strains in advance. Impact of *B. cereus* and cereulide is urgently needed to stop contamination and toxin production, thereby safeguarding the public health. Cereulide is an extremely stable emetic toxin produced by *B. cereus* that is unlikely to be inactivated by food processing and has a high toxicity and its related hazards raise public health concerns. (Yang *et al*., 2023).

Rapid detection of *B. cereus* in food is important to facilitate the application of quality control measures to eradicate *B. cereus* from food and improve diagnosis of food poisoning outbreaks (Ombui *et al*., 2008). Despite the importance of the *B. cereus* group as major foodborne pathogens that may cause diarrheal and/or emetic syndrome(s) (Amor *et al*., 2019), no study in Haryana has been conducted in order to characterize the occurrence of the *B. cereus* group in milk products. As Food, the most important energy source, might be easily contaminated by pathogens if not handled hygienically (Mead *et al*., 1999).

## Materials and methods

### Sample collection

A total of 200 samples of milk products were collected from retail shops in Haryana state (Fig. 1). As Haryana state is divided into two agroclimatic zones, viz. Southwestern (SW) zone and Northeastern (NE) Zone. From each agroclimatic zone, two districts and from each selected district, two tehsil were selected randomly. Thereby, a total of four district (Hisar, Rewari, Karnal, Panchkula) and eight tehsils (Hisar, Hansi, Rewari, Kosli, Karnal, Indri, Panchkula, Kalka) were selected from the Haryana state. Then, from each tehsil five sweet shops selected and from each shop a five milk products sample Peda, Milk Cake, White Burfi, Rajasthani Burfi, Mawa Burfi and others will be collected by simple random sampling. A total 200 samples were collected.

**Fig 1.**
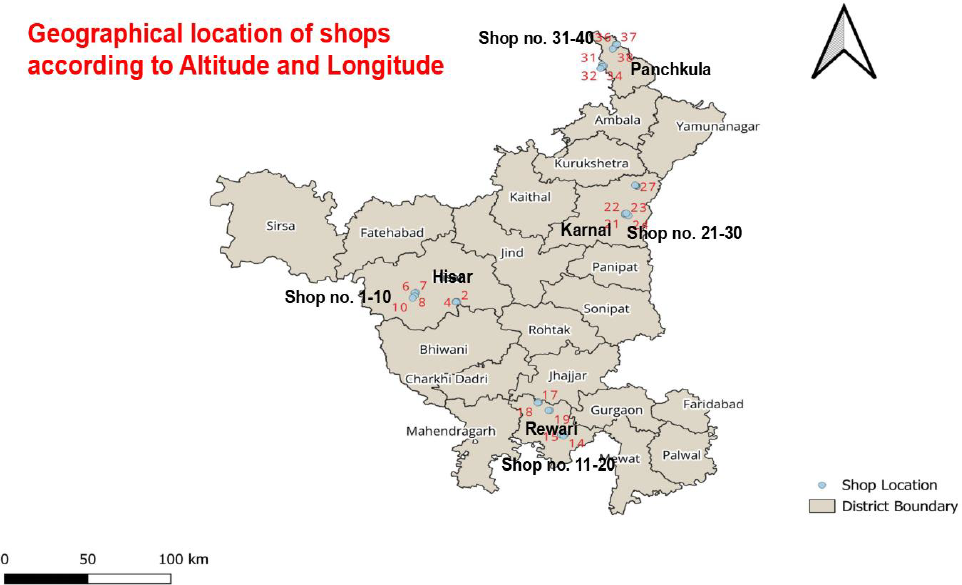
Geographical map showing different sweets shops.

The samples from different places were collected in interval of three months at regular interval. Immediately after collection, the samples were transported to the laboratory in insulated container containing ice gel packs for analyses. All the samples tested in this study were collected between September 2023 and November 2023.

### Isolation and characterization of Bacillus species

Isolation of Bacillus spp. was done as described previously (Public Health England 2018; Tewari *et al*., 2015). A portion (1 g) from the centre of each sample was collected aseptically and was homogenised with 9 ml of brain heart infusion (BHI) broth. This homogenate was enriched by incubating at 37°C for 24 h. 100 mcl of enriched samples were streaked on polymyxin pyruvate egg yolk mannitol bromothymol blue (PEMBA, HiMedia Lab, Mumbai, India) agar plates in duplicate and incubated at 37°C for 24 h (Tewari *et al*., 2015). The fimbriate peacock blue coloured colonies surrounded by a blue zone of egg yolk hydrolysis showing Bacillus like growth characteristics were purified for identification by biochemical tests and carbohydrate utilization reactions (Public Health England 2018; Vos *et al*., 2009). All the chemicals and reagents used in biochemical characterization of Bacillus isolates were purchased from HiMedia laboratories, Mumbai, India. Presumptive identification of *B. cereus* was carried out by colony characteristics, Gram staining (Gram positive) and IMViC test (--++) (Cowan and Steels, 1974). Other biochemical tests that were Nitrate reduction (Positive), Oxidase test (negative), and Catalase test (Positive).

### DNA extraction and amplification reactions

DNA extracted via snapchill method. Inoculate a loopfull of bacterial suspension in 2 ml of BHI broth at 37 °C for 24-48 h till pellet fomation at base. Centrifuse inoculated broth at 8000 rpm for 3 min twice and discard supernatent. Give 2 washings of remaining pellet with 200uL PBS buffer by centrifusing at 14000 rpm for 5 min each washing and discard supernatent. Add 200 uL of Nuclease Free Water (NFW) and vortex well so that pellet can dissolve. Heat the tube in hot plate at 100 °C for 10 min followed by quick chilling at −20 °C for 10 min. After this centrifuse the tube at 14000 rpm for 5 min and take supernatent in sterile DNA tubes. Check concentration of DNA and store at −20°C.

Genes selected in this study shown in table 1. Polymerase chain reaction (PCR) amplifications (Gen eAmp PCR System 9700, LABINDIA, India) were done to reconfirm biochemically characterized Bacillus isolates and to ascertain toxigenic profiles. All primers used for DNA amplification reactions were purchased from Integrated DNA Technologies. Bacillus isolates characterized biochemically were reconfirmed by PCR targeting *16SrRNA* (Gomaa and Momtaz 2006). PCR mixture (25 ul) contained, template DNA (5 ll), GoTaq Green Master Mix (12.5ul, Thermo Fisher Scientific, Mumbai, India), nuclease free water (6.5ul, Thermo Fisher Scientific, Mumbai, India), 0.5ul of forward primer (10 pmol/ll), 0.5 ull of reverse primer (10 pmol/ll). Amplifications were done by using primers (PA-F: 5’-AGAGTTTGATCCTGGCTCAG-3’ and PH-R:3’-AAGGAGGTGATCCAGCCGCA-5’) and amplification parameters as described by Gomaa and Momtaz (2006). DNA amplifications were carried out using PCR conditions; initial denaturation at 95°C for 5 min, followed by 30 cycles of amplification at 95°C for 45s (denaturation), 64°C for 1 min (annealing), 72°C for 1 min (extension) and final extension at 72°C for 5 min.

**Table 1.**
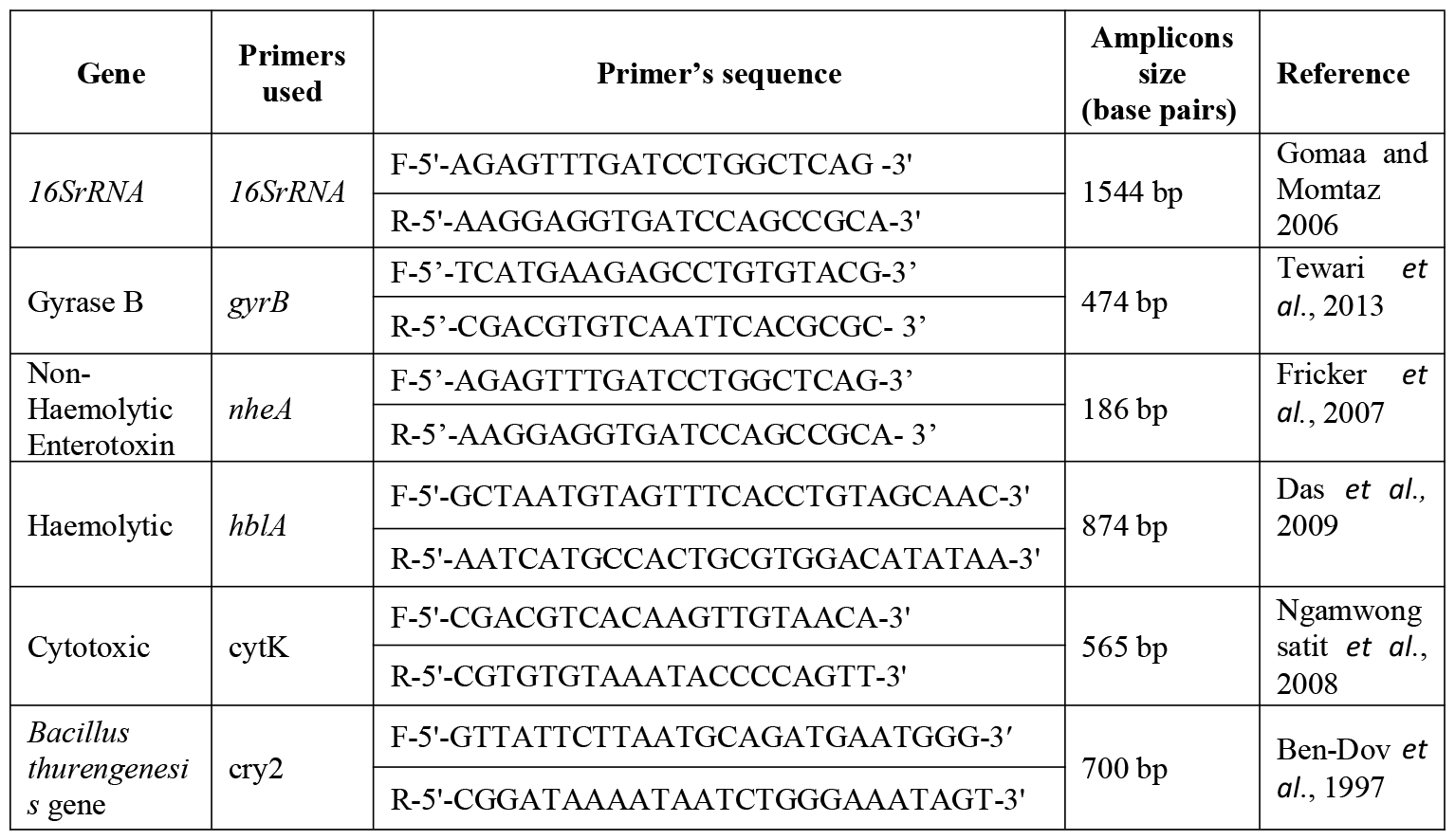
Genes used in research.

PCR was performed to detect the genes encoding gyrase (*gyrB*), haemolytic BL (*hblA*), non haemolytic NHE (*nheA*) and cytotoxic-K (cytK) enterotoxins in Bacillus spp. Isolates. Incidence of Bacillus spp. in khoa based milk products collected from different places of Haryana. Amplification conditions for enterotoxin genes were given in Table 2.

**Table 2.** PCR Conditions for different genes.

| PCR Conditions |  |  |  |  |  |
| --- | --- | --- | --- | --- | --- |
| Gene Designation | Initial denaturation | Amplification (× 35) |  |  | Final extension |
|  |  | Denaturation | Annealing | Extension |  |
| *16SrRNA* | 95 °C for 5 min | 95 °C for 45s | 64 °C for 60s | 72 °C for 60s | 72 °C for 5 min |
| *gyrB* | 95 °C for 5 min | 95 °C for 45s | 64 °C for 60s | 72 °C for 60s | 72 °C for 5 min |
| *nheA* | 95 °C for 5 min | 95 °C for 45s | 62 °C for 60s | 72 °C for 60s | 72 °C for 5 min |
| *hblA* | 95 °C for 5 min | 95 °C for 45s | 70 °C for 60s | 72 °C for 60s | 72 °C for 5 min |
| *cytK* | 95 °C for 5 min | 95 °C for 45s | 62 °C for 60s | 72 °C for 60s | 72 °C for 5 min |
| *cry2* | 95 °C for 5 min | 95 °C for 45s | 60 °C for 60s | 72 °C for 60 | 72 °C for 5 min |

PCR amplicons for 16S rRNA and toxin genes were analyzed by electrophoresis on 1% agarose gels using 100 bp DNA ladders (Thermo Fisher Scientific, Mumbai, India). Agrose gels were stained with ethidium bromide (10 lg/100 ml) and were visualized under UV trans-illuminator (Alphalmager, Alpha Innotech Co., SanLeand, USA).

### Antimicrobial susceptibility testing

Antimicrobial susceptibility testing was performed using standard disc diffusion method on Mueller–Hinton agar (Bauer *et al*., 1966) for fourteen different antibiotics given in table 3.

**Table 3.** List of antibiotics that were used in research.

| Antimicrobials | Disc conc. | Antimicrobials | Disc |
| --- | --- | --- | --- |
| Amikacin (AK) | 30 mcg | Gentamycin (GEN) | 10 mcg |
| Ampicillin (AMP) | 10 mcg | Imipenem (IPM) | 10 mcg |
| Azithromycin | 15 mcg | Levofloxacin (LE) | 5 mcg |
| Cefazolin (CZ) | 30 mcg | Meropenem (MRP) | 10 mcg |
| Ciprofloxacin | 5 mcg | Methicillin (MT) | 5 mcg |
| Colistin (CLR) | 15 mcg | Polymyxin-B (PB) | 300 mcg |
| Enrofloxacin (Ex) | 10 mcg | Penicillin (P) | 10 mcg |

Antibiotic discs were placed on Mueller–Hinton agar plates inoculated with *B. cereus* isolates at 37°C and zones of inhibition were measured after 24h. Antibiotic concentrations of the discs used and antimicrobial susceptibility interpretation criteria were as recommended by the Clinical and Laboratory Standards Institute (CLSI 2013). Isolates resistant to 3 or more unique antimicrobial classes were classified as multi drug resistant (MDR) strains (Waters *et al*., 2011).

### Statistical analysis

Fisher’s exact test was used to determine the association between incidence of *B. cereus* and its enterotoxins with and types of RTE foods tested in this study. Significance was set at a two tailed p value <0.05.

### Results and discussion

*Bacillus cereus* is one of 22 World Health Organization prioritized food borne pathogens for assessing the load of food borne illnesses (Kirk et al., 2015). This investigation determined the burden of *B. cereus* and its enterotoxins in khoa based products (White burfi, Peda, Milk cake, Rajasthani burfi, mawa burfi) from different districts in Haryana. As consumption of these sweets remains higher always due to continues demand. These foods are generally prepared and stored in an uncontrolled environment without following good hygienic and manufacturing practices. High levels of food safety standards should be followed while preparing and selling these products. Of 200 samples tested, Bacillus isolates were recovered from 88 samples with an overall incidence of 44%. PCR amplification of 16S rRNA (1544 bp, Gomaa and Momtaz 2006; Sadashiv and Kaliwal 2014) confirmed all biochemically characterized isolates (94) as Bacillus spp. (Fig. 1). Organji et al. (2015) and Hadithi et al. (2016) had reported higher Bacillus incidence rates of 17.3% and 24.76%, respectively in foods comprising cooked rice, pasteurized milk, yogurt, infant milk powder and white cheese. Incidence of Bacillus was highest in white burfi (50%) followed by Rajasthani burfi (47.5%), milk cake (45%), Mawa burfi (40%), Peda (35%). Tehsil wise prevalance is noted in table 4.

**Table 4.** Positivity of different samples in different tehsils according to *16SrRNA* gene.

| Tehsil | Total positive isolates | Peda | Milk cake | White Burfi | Raj. Burfi | Mawa Burfi |
| --- | --- | --- | --- | --- | --- | --- |
| Hansi | 10 | 1/5 | 3/5 | 2/5 | 3/5 | 1/5 |
| Hisar | 8 | 1/5 | 2/5 | 2/5 | 2/5 | 1/5 |
| Rewari | 9 | 1/5 | 1/5 | 2/5 | 2/5 | 3/5 |
| Kosli | 6 | 1/5 | 1/5 | 1/5 | 1/5 | 2/5 |
| Karnal | 15 | 2/5 | 4/5 | 4/5 | 2/5 | 3/5 |
| Indri | 8 | 2/5 | 1/5 | 3/5 | 2/5 | 0/5 |
| Panchkula | 15 | 2/5 | 3/5 | 3/5 | 4/5 | 3/5 |
| Kalka | 17 | 5/5 | 3/5 | 3/5 | 3/5 | 3/5 |
| **Total** | **88/200** | **15/40** | **18/40** | **20/40** | **19/40** | **16/40** |
|  | **(44%)** | **(35%)** | **(45%)** | **(50%)** | **(47.5%)** | **(40%)** |

*B. cereus* is major food borne pathogen from genus Bacillus, whereas *B. alvei, B. polymyxa and B. firmus* are more associated with food spoilage (Hanlin 1998). All identified *B. cereus* isolates were Gram positive rods. These isolates were catalase, Nitrate reduction, Voges Proskauer and citrate positive and negative for indole, methyl red test. We recorded an overall *B. cereus* incidence of 44% (88/200). Tewari *et al*. (2015) and Fayaz *et al*. (2017) reported 30.9% and 36.7% incidence of *B. cereus* in meat and meat products, respectively. Yusuf *et al*. (2018) reported *B. cereus* in 28.4% samples of milk and milk products.

Abbas *et al*. (2014) reported 16.7% incidence of *B. cereus* in soft cheese. Salem *et al*. (2015) reported 20% samples each of white cheese and Kareesh cheese contaminated with *B. cereus* in Egypt. Iurlina *et al*. (2006) reported 50% incidence of *B. cereus* in Port Salut Argentino cheeses. Singh *et al*. (2015) found 52.9% paneer samples contaminated with *B. cereus*, much higher than recorded in this study. Yusuf *et al*. (2018) found 16% paneer samples contaminated with *B. cereus*. High incidence of *B. cereus* in cheeses and other dairy products can be due to high hydrophobicity of the spores, the low spore surface charge and the spore morphology which results in their strong adhesion to different surfaces and survival during processing of the milk and milk products (Andersson *et al*., 1995; Iurlina *et al*., 2006). All 88 Bacillus isolates were screened by PCR for the presence of enterotoxin genes; *nheA* (186 bp), *hblA* (874 bp) *gyrB* (474 bp) and cytK (565 bp). None of the *B. cereus* isolates carried cry2 gene that is coded by *B. thurengenesis*. Owusu-Kwarteng *et al*. (2017) reported 92% *B. cereus* isolates from dairy farms and traditional dairy food products positive for one or more nhe complex encoding genes. Similarly, Tewari *et al*. (2015) reported 89.7% of *B. cereus* isolates with at least one of the genes of nhe complex. Among nhe complex genes, we observed highest incidence of nheC gene (100%, 19/19) followed by nheB (94.7%, 18/19) and *nheA* (57.9%, 11/19). Abbas *et al*. (2014) reported higher presence of *nheA* (90.3%) compared to nheB (58.1%) and nheC (54.8%) in *B. cereus* isolated from milk and milk products. 57.9% *B. cereus* isolates in this study contained all three nhe genes, 36.8% contained two genes (nheB, nheC) and 5.3% had only one nhe gene i.e. nheC. Using qPCR/MiSeq sequencing data, high incidence of *nheA* (135/145), nheB (113/145) and nheC (145/145) was detected in 147 fermented soybean products contaminated with *B. cereus* (Keisam et al., 2019). In our study, incidence of *B. cereus* cytK was (52.6%, n = 10/19). Earlier studies had reported incidence ranging from 41.4% to 88.0% (Ngamwongsatit *et al*., 2008; Rather *et al*., 2011; Tewari *et al*., 2015; Owusu-Kwarteng *et al*., 2017). *B. cereus* enterotoxigenic genes in an operon (hblCDA and *nheA*BC) can occur independently of each other. Tewari *et al*. (2015) had reported independent occurrence of hbl, nhe and cytK enterotoxin in *B. cereus*.

**Fig. 2.**
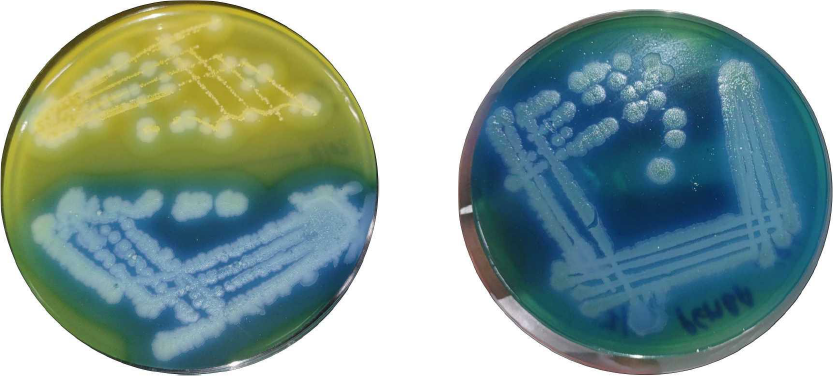
Yellow colour colonies are Negative for *B. cereus* group while Blue colour colonies are positive for *B. cereus* group.

The positivity of different genes in different tehsils of Haryana shown in table 5.

**Table 5.** Positivity of different genes in different tehsils.

| Location | *16SrRNA* positivity | *gyrB* gene positivity | *nheA* gene positivity | *hblA* gene positivity | *CytK* gene positivity |
| --- | --- | --- | --- | --- | --- |
| Hansi ( n = 25 ) | 10 (40%) | 2 (8%) | 8 (32%) |  | 1 (4%) |
| Hisar ( n = 25 ) | 8 (32%) | 1 (4%) | 6 (24%) |  | 1 (4%) |
| Rewari ( n = 25 ) | 9 (36%) | 0 | 5 (20%) |  | 0 |
| Kosli ( n = 25 ) | 6 (24%) | 0 | 3 (12%) |  | 0 |
| Karnal ( n = 25 ) | 15 (60%) | 4 (16%) | 9 (36%) |  | 4 (16%) |
| Indri ( n = 25 ) | 8 (32%) | 1 (4%) | 2 (8%) |  | 1 (4%) |
| Panchkula ( n = 25 ) | 15 (60%) | 3 (12%) | 12 (48%) | 1 (4%) | 0 |
| Kalka ( n = 25 ) | 17 (68%) | 2 (8%) | 5 (20%) |  | 2 (8%) |
| **Total ( 200 )** | 88/200<br>(44%) | 13/200<br>(6.5%) | 50/200<br>(25%) | 1/200<br>(0.5%) | 9/200<br>(4.5%) |

**Fig. 3.**
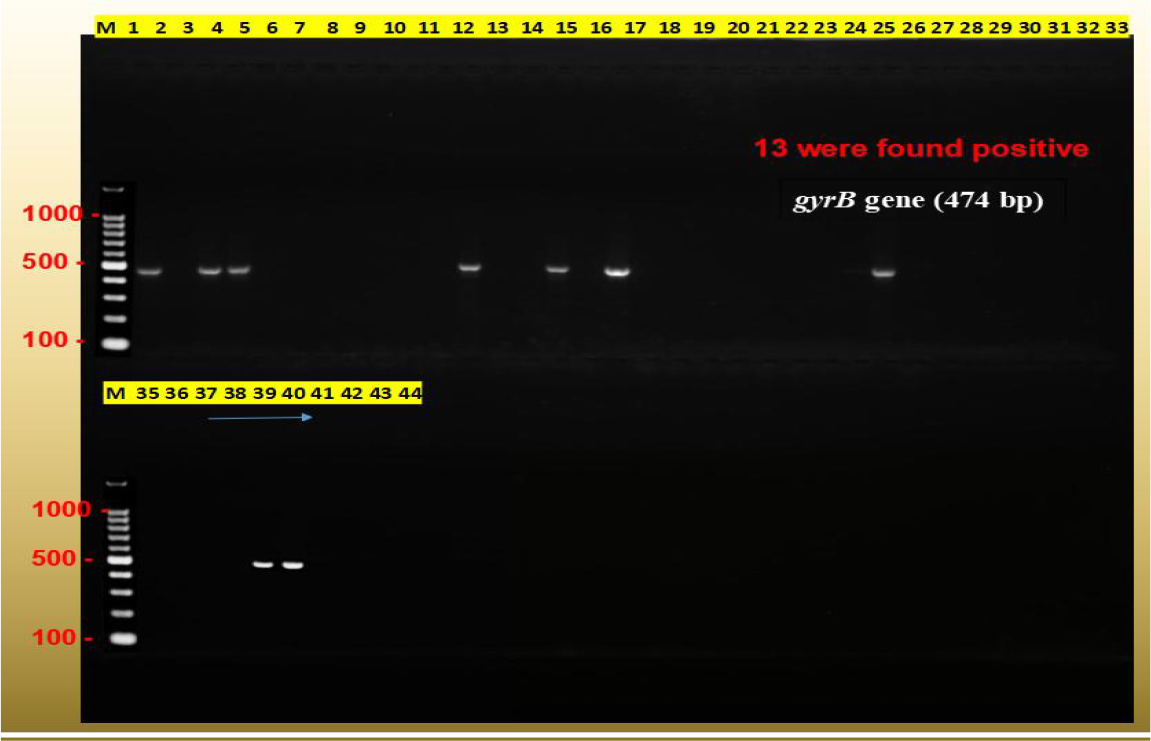
Gel electrophoresis showing *gyrB* (474 bp) gene of *B. cereus*, amplified from *B. cereus* isolates. Lane M– 100bp DNA ladder; Lane 1– Positive control; Lane 2– Negative control; Lanes 3-44 – *B. cereus* isolates

**Fig. 4.**
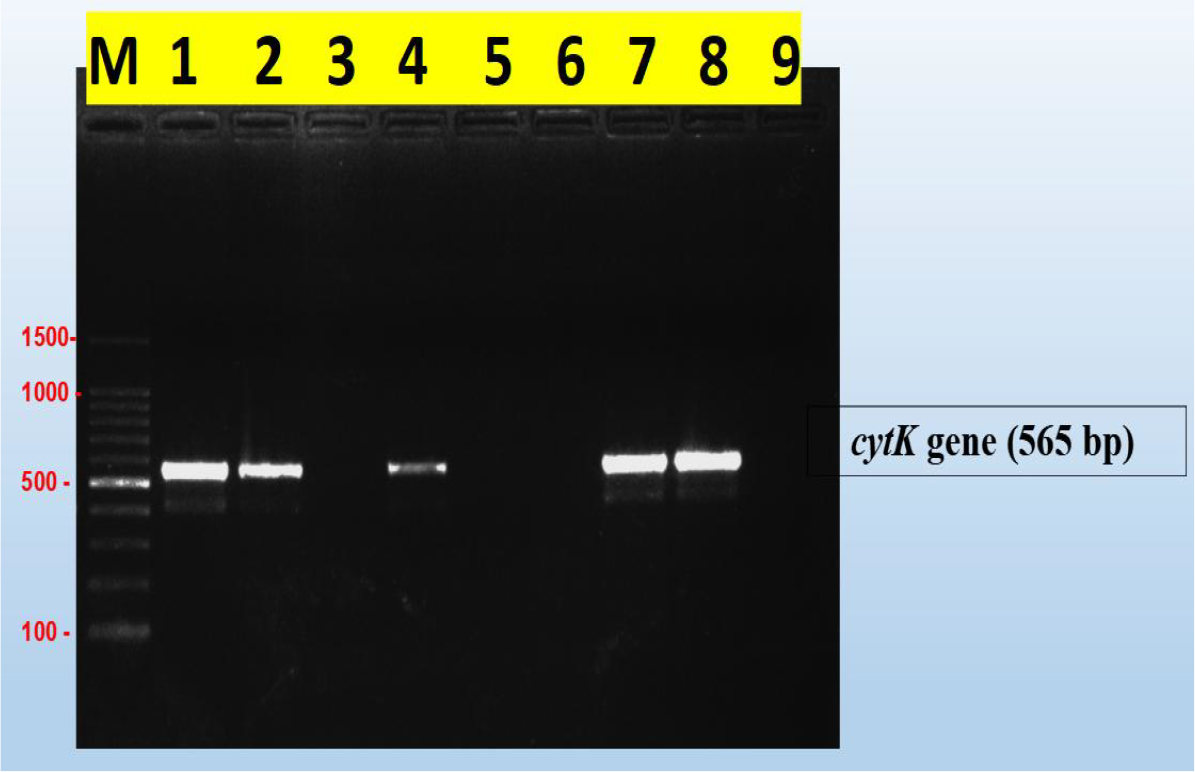
Gel electrophoresis showing cytK (565 bp) gene of *B. cereus*, amplified from *B. cereus* isolates. Lane M– 100bp DNA ladder; Lanes 1-7– *B. cereus* isolates; Lane 8 – Positive control; Lane 9 – Negative control

All the 88 isolates were subjected to Antibiotic susceptibility testing against 14 different antibiotics of different classes. Antibiotics were selected according to CLSI guidelines list (CLSI 2013). The Antibiotic susceptibility pattern of the isolates is given in table 6. Maximum sensitivity of was observed for Amikacin, Gentamycin, Levofloxacin, Meropenem (100% each), followed by Azithromycin (71.6%), Colistin (54.5%), imipenem (43.2%). High antibiotic resistance was observed for Ampicillin, Cefazolin, Methicillin, Polymyxin-B, Penicillin (100% each) shown in table 6. All isolates were multiple drug resistant (MDR) as resistant for three or more classes of antibiotics was present.

**Table 6.**
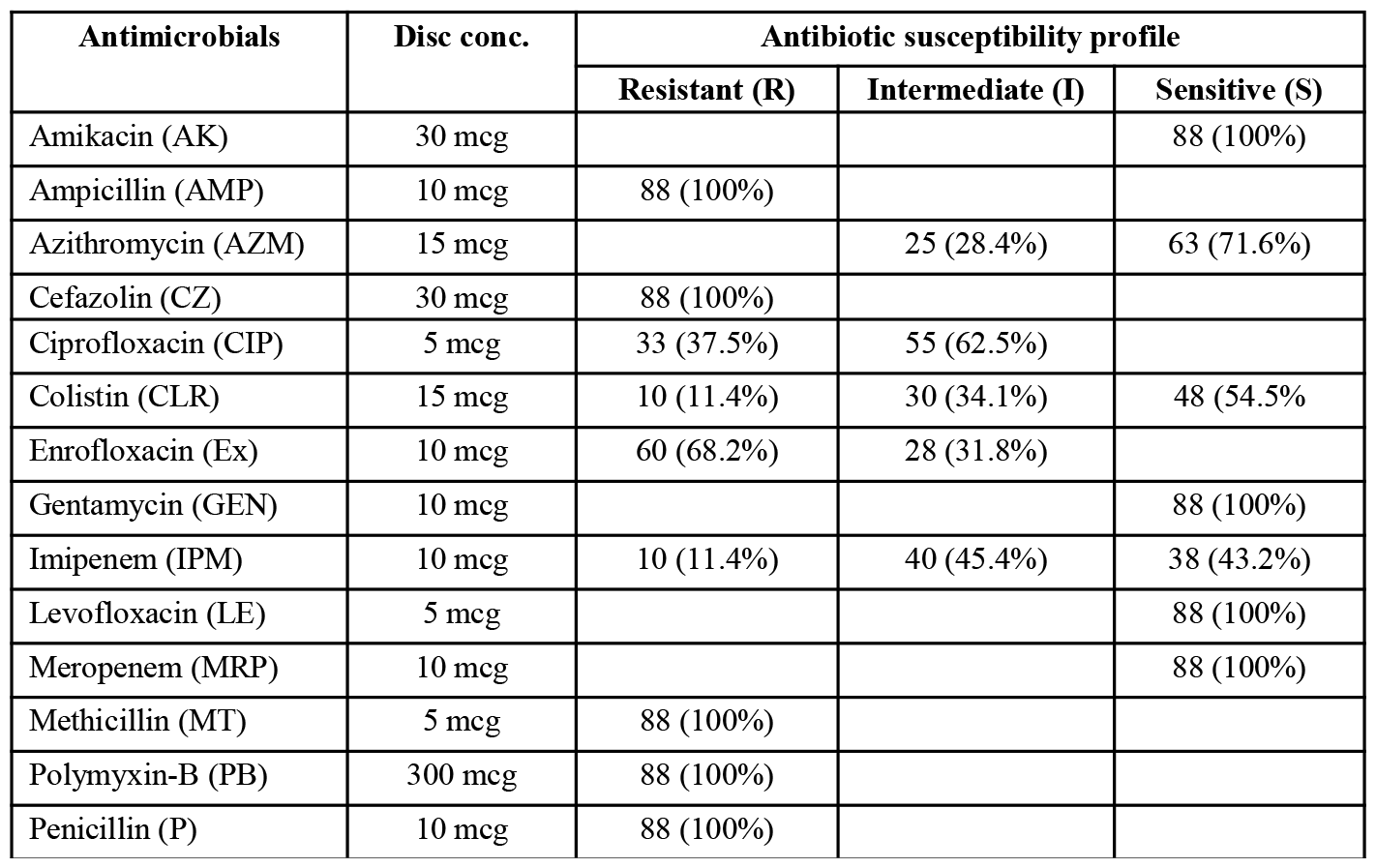
Antibiotic susceptibility profile of *B. cereus* isolates (n=88)

All the *B. cereus* isolates were multi-drug resistant. Each of those was resistant to at least five different antibiotics used. Most of the isolates were resistant to Ampicillin, Cefazolin, Methicillin, Polymyxin-B, Penicillin. In another study all the 48 isolates of *B. cereus* from legume-based fermented food products were resistant against this higher concentration of polymyxin B (Roy *et al*., 2007). An investigation on antibiotic-resistance profiles of *B. cereus* isolates from different food products in Morocco showed that the isolates were resistant to ampicillin, tetracycline and penicillin, but susceptible to chloramphenicol and erythromycin (Merzougui *et al*., 2014). Çürek *et al*. (2023) found that all *B. cereus* isolates from spices were susceptible to linezolid and vancomycin, while 35.42% showed resistance to erythromycin. Multi-drug resistance was detected in 8.3% of the isolates. Fiedler *et al*. (2019) reported that *B. cereus* group strains isolated from fresh vegetables showed resistance against β-lactam antibiotics such as penicillin G and cefotaxime, as well as amoxicillin/clavulanic acid combination and ampicillin. Most strains were susceptible to ciprofloxacin, chloramphenicol, amikacin, imipenem, erythromycin, gentamicin, tetracycline, and trimethoprim/sulfamethoxazole combination. Meena *et al*. (2019) observed that all the isolates from milk and milk products had 100% sensitivity to amikacin and 100% resistance to ampicillin, methicillin, and penicillin-G. Anonymous (2024) noted that strains were completely resistant to penicillin, amoxicillin-clavulanic acid, cefixime, ceftriaxone, vancomycin, and co-trimoxazole, with a species-specific trend in both phenotypic and genotypic resistance patterns. Anonymous (2020) indicated that most *B. cereus* isolates were resistant to ampicillin, penicillin, cefepime, cephalothin, and oxacillin, and were susceptible to gentamicin, chloramphenicol, imipenem, tetracycline, ciprofloxacin, trimethoprim-sulfamethoxazole, erythromycin, kanamycin, and cefotetan. Kumari & Sarkar (2014) found that *B. cereus* isolates from dairy products were resistant to multiple antibiotics, including penicillin, ampicillin, and vancomycin, highlighting the risk of antibiotic-resistant strains in food products. Tallent *et al*. (2012) demonstrated that *B. cereus* isolates from food matrices exhibited resistance to various antibiotics, emphasizing the need for improved detection and control measures. Liu *et al*. (2020) reviewed the antibiotic resistance profiles of *B. cereus* in dairy products, showing a significant number of isolates resistant to common antibiotics. Springer *et al*. (2020) provided an overview of the antibiotic susceptibility of *B. cereus* clinical isolates, with many showing resistance to β-lactam antibiotics. FIL-IDF (2016) discussed the importance of monitoring antibiotic resistance in *B. cereus* strains from raw milk to ensure food safety.

### Confirmation of *B. cereus* from *B. thurengensis*

As *B. cereus* very similar to *B. thurengenesis* so for differentiation it 2 method used.

#### 1. By staining

By using Fadel et al. (1988) a rapid and simple method for staining of crystal protein of B. thurengenesis that differentiate it from B. cereus. In this method samples were prepared for microscopic examination as follows: smears of bacteria were dipped into a small container containing Coomassie brilliant blue solution (0.25 % Coomassie brilliant blue, 50% ethanol, and 7% acetic acid) for 3 min, washed with tap water, dried, and observed under a light microscope.

#### 2 By Molecular Method :-

*B. thurengenesis* specific gene cry2 was targeted to rule out the isolates as *B. thurengenesis* from the isolates recovered in the present study.

Further sequencing of some isolates was done and a phylogenic tree was made to study the ancestor origin of the *B. cereus* that was isolated. Upto 99 % similarity found in sequences of isolates from other strains found in India and abroad.

## Conclusion

In conclusion, this comprehensive research sheds light on the prevalence, characteristics, and antibiotic resistance patterns of *B. cereus* in milk products across Haryana, India. Through meticulous sampling and analysis, a total of 200 samples from eight tehsils spanning two agroclimatic zones were examined. The isolation and identification process, involving pre-enrichment, streaking on selective media, and biochemical tests, led to the confirmation of 88 positive *B. cereus* isolates, all of which were further distinguished from *Bacillus thuringiensis* via cry2 gene.

Moreover, the analysis of toxic genes revealed the presence of *gyrB*, cytk, *hblA*, and *nheA* genes, indicating the potential virulence of the isolated strains. Geographical variation in prevalence was evident, with higher rates observed in certain tehsils, notably Kalka and Panchkula. Specifically, prevalence ranged from 32% to 68% across the sampled tehsils.

Of significant concern is the antibiotic resistance observed among the isolates, with resistance detected against multiple classes of antibiotics. Notably, all isolates exhibited resistance to Ampicillin, Cefazolin, Methicillin, Polymyxin-B, and Penicillin. Conversely, maximum sensitivity was observed for Amikacin, Gentamycin, Levofloxacin, and Meropenem.

These findings underscore the pressing need for stringent food safety measures and surveillance protocols in dairy production and distribution. Effective interventions are imperative to mitigate the risk of *B. cereus* contamination and combat antibiotic resistance in milk products. By providing detailed insights and data, this research informs evidence-based strategies to ensure the safety and quality of dairy products, thus safeguarding public health in the region. Continued research and monitoring efforts are essential to address emerging challenges and uphold food safety standards effectively.

## Bibliography

Abbas, B.A., Khudor, M.H., & Saeed, B.M.S. (2014). Detection of hbl, nhe and bceT toxin genes in Bacillus cereus isolates by multiplex PCR. International Journal Current Microbiology and Applied Sciences, 3: 1009–1016

Altaf, M. S., Hussain, S. A., Ahmad, C. R., Willayat, M. M., Imtiyaz, A. H., & Bhat, S. A. (2015). A study on the Prevalence of B. cereus Emetic Strains in Raw Milk in and around Srinagar City of J and K. Biomedical and Pharmacology Journal, 4(1): 181–188

Amor, M. G., Jan, S., Baron, F., Grosset, N., Culot, A., Gdoura, R., Gautier, M., & Techer, C. (2019). Toxigenic potential and antimicrobial susceptibility of B. cereus group bacteria isolated from Tunisian foodstuffs. BMC Microbiology, 19(1): 1–12

Anderson Borge, G. I., Skeie, M., Sørhaug, T., Langsrud, T., & Granum, P. E. (2001). Growth and toxin profiles of B. cereus isolated from different food sources. International Journal of Food Microbiology, 69: 237–246

Andersson A, Ronner U, Granum PE (1995) What problems does the food industry have with the spore-forming pathogens Bacillus cereus and Clostridium perfringens? International Journal of Food Microbiology, 28: 145–155

Ankolekar, C., Rahmati, T., & Labbé, R. G. (2009). Detection of toxigenic B. cereus and Bacillus thuringiensis spores in U.S. rice. International Journal of Food Microbiology, 128: 460–466

Anonymous. (2020). Prevalence, molecular characterization, and antibiotic susceptibility of B. cereus isolates from dairy products. Journal of Dairy Science, 103(8): 6961–69713

Anonymous. (2024). Emergence of multidrug-resistant Bacillus spp. derived from animal feed. BMC Microbiology, 24(1): 315

Bacillus Food Poisoning. Cambridge City Council, (https://www.cambridge.gov.uk/sites/www.cambridge.gov.uk/files/docs/Bacillus%20food%20poisoning.pdf)

Bauer, A. W., Kirby, W. M., Sherris, J. C., & Turck, M. (1966). Antibiotic susceptibility testing by a standardized single disk method. American Journal of Clinical Pathology, 45(4): 493–496

Ben-Dov, E., Zaritsky, A., Dahan, E., Barak, Z. E., Sinai, R., & Manasherob, R., (1997). Extended screening by PCR for seven cry-group genes from field-collected strains of Bacillus thuringiensis. Applied and Environmental Microbiology, 63(12): 4883–4890

Bratcher, D. F., Long, S., Prober, C., & Fischer, M. (2017). Bacillus species (Anthrax). In Principles and Practice of Pediatric Infectious Diseases (5th ed., pp. 770–773). Elsevier sci.: Philadelphia

CDC. (2011). Estimates: Findings-CDC estimates of foodborne illness in the United States. Retrieved from http://www.cdc.gov/foodborneburden/2011-foodborneestimates.html

Chang, Y., Xie, Q., Yang, J., Ma, L., & Feng, H. (2021). The prevalence and characterization of B. cereus isolated from raw and pasteurized buffalo milk in southwestern China. Journal of Dairy Science, 104(4): 3980–3989

Chen, L., Daniel, R.M. and Coolbear, T. (2003) Detection and impact of protease and lipase activities in milk and milk powders. International Dairy Journal, 13: 255–275

CLSI (2013) Performance standards for antimicrobial susceptibility testing. Twenty-third informational supplement. CLSI document M100-S23. Clinical and Laboratory Standards Institute, Wayne

Cowan, S. T., & Steel, K. J. (1974). Cowan and Steel’s Manual for the Identification of Medical Bacteria (3rd ed.). Cambridge University Press

Çürek, S., Geniş, B., Tuncer, B. Ö., & Tuncer, Y. (2023). Prevalence, Toxin Genes, and Antibiotic Resistance Profiles of B. cereus Isolates from Spices in Antalya and Isparta Provinces in Türkiye. Indian Journal of Microbiology, 63: 549–5611

Das, S., Surendran, P.K., & Thampuran N. (2009). PCR-based detection of enterotoxigenic isolates of B. cereus from tropical seafood. Indian J Med Res. 129: 316–320

Fayaz S, Badroo GA, Ahmad A, Rasool U, Mustafa R, Mudasir M (2017) Molecular characterization of enterotoxigenic Bacillus cereus species isolated from meat using conventional PCR and multiplex PCR. International Journal Current Microbiology and Applied Sciences, 6: 324–328

Fiedler, G., Schneider, C., Igbinosa, E. O., Kabisch, J., Brinks, E., Becker, B., Stoll, D. A., Cho, G. S., Huch, M. & Franz, C. M. (2019). Antibiotics resistance and toxin profiles of Bacillus cereus-group isolates from fresh vegetables from German retail markets. BMC microbiology, 19(1): 1–13

FIL-IDF (2016). B. cereus in raw milk: Hygiene criteria and control measures. International Dairy Federation Bulletin, 491: 1–158

Fricker, M., Messelhäusser, U., Busch, U., Scherer, S., & Ehling-Schulz, M. (2007). Diagnostic real time PCR assay for the detection of emetic B. cereus strains in foods and recent food borne outbreaks. Applied and Environmental Microbiology, 73: 1892–1898

Gomaa, O. M., & Momtaz OA (2006). 16SrRNA characterization of a Bacillus isolate and its tolerance profile after subsequent subculturing. Arab J Biotechnol 10: 107–116

Granum, P. E., & Lund, T. (1997). B. cereus and its food poisoning toxins. FEMS Microbiol. Lett., 157: 223–228

Haddaji, N., Chakroun, I., Fdhila, K., Smati, H., Bakhrouf, A., & Mzoughi, R. (2022). Pathogenic impacts of B. cereus strains on Crassostrea gigas. Journal of Ecology, 19(2): 1–8

Hanlin, J.H. (1998) Spoilage of acidic products by Bacillus species. Dairy Food Environment Sanitation, 18: 655–659

Iurlina, M.O., Saiz, A.I., Fuselli, S.R. & Fritz, R. (2006). Prevalence of Bacillus spp. in different food products collected in Argentina. LWT, 39: 105–110

Jessberger, N., Kranzler, M., Riol, C. D., Schwenk, V., Buchacher, T., Dietrich, R., Ehling-Schulz, M., & Martlbauer, E. (2019). Assessing the toxic potential of enteropathogenic B. cereus. Journal of Ecology, 84: 103276

Johnson, D. A., Aulicino, P. L., & Newby, J. G. (1984). B. cereus induced myonecrosis. Journal of Trauma, 24(3): 267–270

Keisam, S., Tuikhar, N., Ahmed, G., & Jeyaram, K. (2019). Toxigenic and pathogenic potential of enteric bacterial pathogens prevalent in the traditional fermented foods marketed in the Northeast region of India. International Journal of Food Microbiology, 296, 21–28

Khudor, H. M., Abbas, A. B., & Saeed, M. S. B. (2012). Molecular detection of enterotoxin (cyt k) gene and antimicrobial susceptibility of B. cereus isolates from milk and milk products.Basra Journal of Veterinary Research, 11(1): 164–173

Kramer, J. M., & Gilbert, R. J. (1989). B. cereus and other Bacillus species. In Foodborne Bacterial Pathogens (pp. 21–70)

Kumari, S., & Sarkar, P. K. (2014). Antibiogram of isolates of B. cereus group using different antibiotics. Journal of Food Safety, 34(2): 123–1309

Liu, X.-Y., Hu, Q., Xu, F., Ding, S.-Y., & Zhu, K. (2020). Characterization of B. cereus in Dairy Products in China. Toxins, 12(7): 4545

Mead, P. S., Slutsker, L., Dietz, V., McCaig, L. F., Bresee, J. S., Shapiro, C., Griffin, P. M., & Tauxe, R. V. (1999). Food-related illness and death in the United States. Emerging Infectious Diseases, 5: 607–625

Meena, S. C., Gaurav, A., Shekhawat, S. S., Joseph, B., Kumar, H., & Kumar, N. (2019). Isolation and Identification of B. cereus from Milk and Milk Products in Udaipur, Rajasthan, India. International Journal of Current Microbiology and Applied Sciences, 8(9): 2783–2787

Merzougui, S., Lkhider, M., Grosset, N., Gautier, M., & Cohen, N. (2014). Prevalence, PFGE typing, and antibiotic resistance of B. cereus group isolated from food in Morocco. Foodborne Pathogens and Disease, 11: 145–149

NDDB (National Dairy Development Board). 2023. Milk production in India. Available at: http://www.nddb.org/information/stats/milkprodindia

Ngamwongsatit, P., Buasri, W., Pianariyanon, P., Pulsrikam, C., Ohba, M., Assavanig, A., & Panbangred, W. (2008). Broad distribution of enterotoxin genes (hblCDA, nheABC, cytK, and entFM) among Bacillus thuringiensis and B. cereus as shown by novel primers. International Journal of Food Microbiology, 121: 352–356. 10.1016/j.ijfoodmicro.2007.11.013. PMid: 18068844.

Nickerson, S.C. 1995. Milk production: Factors affecting milk composition. In: Milk quality. Springer, US, pp. 3–24

Ombui, J.N., Gitahi, J.N., Gicheru, M.M. 2008. Direct detection of B. cereus enterotoxin genes in food by multiplex Polymerase Chain Reaction. International Journal of Integrative Biology, 2(3): 172–181

Owusu-Kwarteng, J., Wuni, A., Akabanda, F., Debrah, K.T., & Jespersen, L. 2017. Prevalence, virulence factor genes and antibiotic resistance of B. cereus sensu lato isolated from dairy farms and traditional dairy products. BMC Microbiology, 17: 65

Prod’hom, G., Bille, J., Cohen, J., Powderly, W.G., Opal, S.M. 2017. Aerobic Gram-positive Bacilli. In Infectious Diseases, Elsevier Science, 4: 1537–1552.e2

Public Health England (2018) UK standards for microbiology investigations. Identification of Bacillus species. Standards Unit, Microbiology Services, Public Health England. Bacteriologyidentification 3(1): pp 1–27. https://assets.publishing.service.gov.uk/government/uploads/system/uploads/attachment_data/file/697260/ID_9i3.1.pdf

Rather, M.A., Aulak, R.S., Gill, J.P.S., Rao, T.S., & Hassan, M.N. (2011) Direct detection of Bacillus cereus and its enterotoxigenic genes in meat and meat products by polymerase chain reaction. J Adv Vet Res, 1: 99–104

Reyes, J.F., Cagnasso, M.A., Corser, P.I., D’Pool, G., Urdaneta, A.G., Leal, K.V. (2001) Antimicrobial resistance of Bacillus isolated from raw milk. Universidal-del-Zulia, 11(6): 479–484

Roy, A., Moktan, B., & Sarkar, P.K. (2007) Characteristics of B. cereus isolates from legume-based Indian fermented foods. Food Control, 18(12): 1555–1564

Salem, N.A., Jakee, J.E., Nasef, S.A. & Badr, H. (2015) Prevalence of Bacillus cereus in milk and milk products. Animal Health Resource Journal, 3:168–172

Schoeni, J.L., & Wong, A.C.L. (2005) B. cereus - Food Poisoning and Its Toxins. Journal of Food Protection, 68(3): 636–648

Singh, V.K., Shukla, S., & Chaturvedi, A. (2015) Study the incidence of Bacillus cereus isolates from dairy foods. Pharma Innovation Journal, 3: 41–43

Springer, S., Selb, R., & Eisenberg, T. (2020) Prevalence, molecular characterization, and antibiotic susceptibility of B. cereus in dairy products in Germany. Journal of Dairy Science, 103(8): 6961–6971

Suthar, A.P., Kumar, R., Savalia, C.V., Patel, N.M., & Kalyani, I.H. (2019) Toxigenic Profiling of Enterotoxin-Producing B. cereus Isolated from Marketed Raw Chicken Meat and Human Subjects by Triplex and Multiplex PCR. Journal of Animal Research, 9(6): 821–829

Svensson, B., Monthan, A., Guinebretiere, M.H., Nguyen, C., Christiansson, A. (2007) Toxin production potential and the detection of toxin genes among strains of the B. cereus group isolated along the dairy production chain. International Dairy Journal, 17(10): 1201–1208

Tallent, M.S., Kotewicz, M.K., Strain, A.E., & Bennett, W.R. (2012) Efficient Isolation and Identification of B. cereus Group. Journal of AOAC International, 95(2): 446–451

Tewari, A., Singh, S.P., & Singh, R. (2013) Incidence and enterotoxigenic profile of B. cereus in meat and meat products of Uttarakhand, India. Journal of Food Science and Technology. DOI 10.1007/s13197-013-8461162-0.

Tewari, A., Singh, S.P., & Singh, R. (2015) Incidence and enterotoxigenic profile of Bacillus cereus in meat and meat products of Uttarakhand, India. J Food Sci Technol, 52:1796–1801

Vos, P.D., Garrity, G.M., Jones, D., Krieg, N.R., Ludwig, W., Rainey, F.A., Schleifer, K.H., & William, B.W. (2009) Bergey’s manual of systematic bacteriology, 3(2): The Firmicutes. Springer, Dordrecht

Waters, A.E., Contente-Cuomo, T., Buchhagen, J., Liu, C.M., Watson, L., Pearce, K., Foster, J.T., Bowers, J., Driebe, E.M., Engelthaler, D.M., Keim, P.S., & Price, L.B. (2011) Multidrug-resistant Staphylococcus aureus in US meat and poultry. Clinical Infectious Diseases, 52:1227–1230

WHO (2007) Food safety and food-borne illness. Fact sheet no. 237. World Health Organization, Geneva, Switzerland.

Yang, S., Wang, Y., Liu, Y., Jia, K., Zhang, Z., & Dong, Q. (2023) Cereulide and Emetic B. cereus: Characterizations, Impacts and Public Precautions. Foods, 12(4): 833

Yu, P., Yu, S., Wang, J., Guo, H., Zhang, Y., Liao, X., Zhang, J., Wu, S., Gu, Q., & Xue, L. (2019) B. cereus isolated from vegetables in China: Incidence, genetic diversity, virulence genes, and antimicrobial resistance. Frontiers in Microbiology, 10: 948

Yu, S., Yu, P., Wang, J., Li, C., Guo, H., Liu, C., Kong, L., Yu, L., Wu, S., & Lei, T. (2020) A study on prevalence and characterization of B. cereus in ready-to-eat foods in China. Frontiers in Microbiology, 10: 3043

Yusuf, U., Kotwal, S.K., Gupta, S., & Ahmed, T. (2018) Identification and antibiogram pattern of Bacillus cereus from the milk and milk products in and around Jammu region. Vet World, 11: 186–191

Yusuf, U., Kotwal, S.K., Gupta, S., & Ahmed, T. (2018). Identification and antibiogram pattern of B. cereus from the milk and milk products in and around Jammu region. Veterinary World, 11: 186–191

